# Peripheral Nociceptor Activity and Placebo Hypoalgesia: A Proof-of-Concept Study

**DOI:** 10.64898/2026.03.05.708556

**Authors:** Lisa Marie Garcia, Andrea Fiebig, Barbara Namer, Susanne Becker

## Abstract

Placebo effects illustrate clearly how cognitive states can influence physiological processes. While central mechanisms of placebo hypoalgesia, i.e. reduce sensitivity to painful stimuli due to expectations, are well established, it remains unclear whether such expectations modulate also peripheral activity. In this proof-of-principle study, we combined a well-established placebo paradigm with microneurography to directly assess responses of single nerve fibers in awake humans. Six healthy participants (18 fibers) underwent each a placebo and a control condition in a randomized crossover design. Placebo hypoalgesia was induced using verbal suggestion and classical conditioning. Manipulation checks confirmed that participants expected hypoalgesic effects of a placebo gel after conditioning. At the peripheral level, data on the recovery cycle of the nerve fibers indicated changes in the excitability of sleeping nociceptors (CMi) in the placebo compared to control condition, namely with enhanced and longer-lasting early subnormality. While other microneurographic outcome measures showed no consistent placebo effects, rather strong time-dependent reductions in fiber responsiveness independent of the experimental conditions were observed. These findings provide preliminary evidence that modulation of peripheral nociceptor activity by cognitive states, specifically expectations, is possible. Such modulation of peripheral activity appears to be fiber-type specific. Time effects have been neglected in microneurographic protocols. The pronounced time effects found here highlight the need for optimized such protocols.

## INTRODUCTION

Placebo effects demonstrate how strongly cognitive processes can influence bodily functions. Such placebo effects are particularly well studied in pain, where placebo hypoalgesia, i.e. reduce sensitivity to painful stimuli due to expectations, is attributed to central mechanisms. In particular, such expectations are assumed to activate descending pathways, which inhibit nociceptive signal transmission at the level of the spinal cord [1]. However, pain perception depends on the functioning of peripheral sensory neurons. In fact, central nociceptive processing relies on input from peripheral nociceptive fibers, whose excitability can be modulated [2]. But, the potential modulation of peripheral nociceptive processing by placebo effects remains unresolved, likely due to the lack of a describable efferent pathway. Though, presynaptic mechanisms related to central sensitization have been proposed which might mediate such effects [3]. Moreover, upregulation of sympathetic activity can increase peripheral nociceptive responses [4]. Some few findings support this idea that expectations may affect peripheral nociception. For example, hypnotic suggestion has been shown to reduce histamine-induced axon reflex erythema, which is mediated by peripheral C-fibers by the subgroup of mechano-insensitive nociceptors in humans [5]. Supporting the link of the sympathetic system and peripheral nociception, nociceptors get spontaneously active under neuropathic conditions when additionally hypoxic conditions are added [6], potentially caused by sympathetic vasoconstriction leading to local hypoxia in the skin.

Here, we combined a well-established placebo paradigm with microneurographic recordings. Microneurography is a specialized neurophysiological technique allowing recordings of activity from single nerve-fibers in awake humans. The placebo paradigm induces expectations of pain relief by a gel (placebo) applied to the skin, which had unbeknownst to the participants no active ingredients. We hypothesized that such pain relief expectations induce placebo hypoalgesia resulting in the modulation of peripheral nociceptor excitability.

## RESULTS AND DISCUSSION

Confirming a successful induction of pain relief expectations, participants rated the effectiveness of the gel (placebo) as high (rating scale 0-10; *M* = 7.58, standard error (*SEM)* = 0.60). Eighteen recorded C-fibers were classified into mechano-sensitive (CM, *n* = 11) and mechano-insensitive C-fibers (CMi, *n* = 7). Due to technical constraints, not all microneurographic protocols could be analyzed for each fiber. Evidence for placebo effect was found in the recovery cycle of CMi-fibers (analyzed in 8 CM and 3 CMi-fibers). All CMi-fibers exhibited a larger latency shift at short interpulse intervals under placebo compared to control, characterized by a more pronounced early subnormal phase and reduced supernormality (Fig. 1A), indicating reduced excitability following prior activation. CM-fibers showed no placebo-consistent effects, although time-dependent changes were observed with a less pronounced early subnormal phase during the 2nd compared to the 1st assessment (Fig. 1B). These findings suggest potential fiber-type specific peripheral placebo effects, which is notable given that CMi-fibers are the main human chemical irritant-nociceptors and involved in inflammatory and neuropathic pain [7].

**Figure 1.**
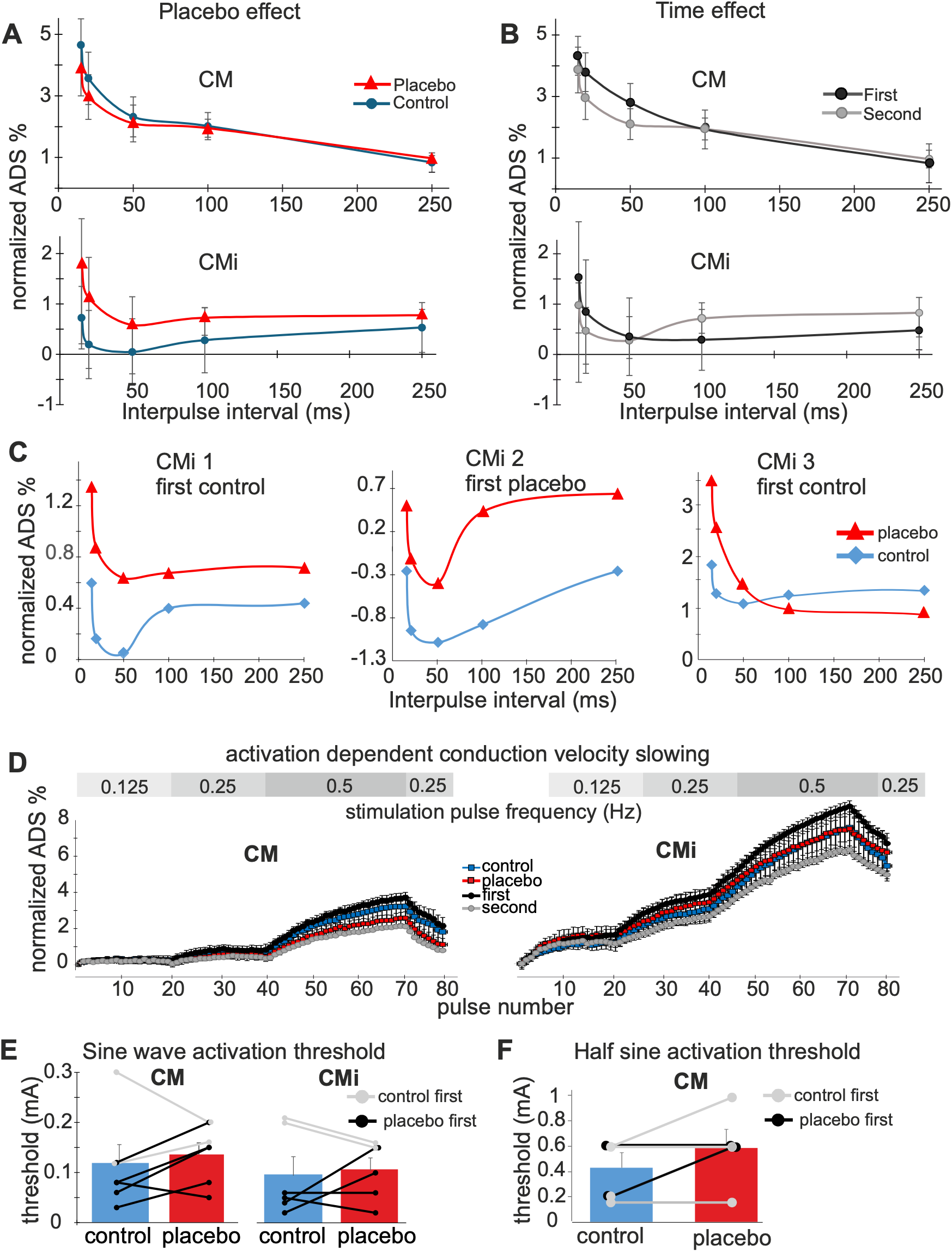
Microneurography results. (A-C). Placebo- and time-effects in the recovery cycle shown as changes in response latency (mean±SEM) due to different interstimulus intervals (ISI) of electrical pulses (activity-dependent conduction velocity slowing (ADS)). **(A)** Shows a placebo effect only in CMi-fibers (*N* = 3). **(B)** Shows a time effect only in CM-fibers (*N* = 8). **(C)** Specimen of the three CMi-fibers of (A). **(D)** Mean ADS (± SEM) in response to low-frequency rising electrical stimulation in CM-fibers (left; *N* = 6; placebo first in 2 fibers) and CMi-fibers (right; *N* = 4; placebo first in 1 fiber). **(E-F)** Single-fiber activation thresholds (mean±SEM) during **(E)** sinusoidal stimulation in CM-fibers (left) and CMi-fibers (right) and **(F)** half-sinusoidal stimulation in CM-fibers (*N* = 5), fibers with placebo first are shown in grey.

Across the other microneurographic measures, placebo effects were inconsistent, but time-dependent changes were prominent. Latency changes during fiber-type identification were consistently smaller during the 2nd compared to the 1st assessment across CM- and CMi-fibers (Fig. 1C), indicating reduced fiber responsiveness over time.

Similarly, thresholds for sinusoidal stimulation increased from the 1st (*M* = 0.09 mA, *SEM* = 0.02 mA) to the 2nd assessment (*M* = 0.14 mA, *SEM* = 0.03 mA) across 13 fibers. Only three fibers showed lower threshold during the 2nd assessment (Fig. 1D). These findings point to a prominent time-dependent modulation of C-fiber excitability that might have outweighed placebo effects. Such time effects might have occurred due to long leg immobilization with compression of the proximal sciatic nerve and reduced blood flow due to sitting still. The present results underscore the need to carefully account for time effects in microneurography studies, which have been neglected.

No time effects were observed for half-sinusoidal stimulation threshold (1st: *M* = 0.50 mA, *SEM* = 0.10 mA; 2nd: *M* = 0.50, *SEM* = 0.16 mA). Instead, these thresholds showed a pattern consistent with placebo hypoalgesia in CM-fibers, with higher mean thresholds under placebo (*M* = 0.58 mA, *SEM* = 0.14 mA) compared to control (*M* = 0.42 mA, *SEM* = 0.11 mA), although these effects were inconsistent across fibers (Fig. 1F). The threshold for the one CMi-fiber was comparable across conditions (placebo: 0.10 mA; control: 0.12 mA).

Pain ratings showed patterns consistent with microneurography results. During sinusoidal stimulation, ratings were substantially lower during the 2nd (*M* = 28.93, *SEM* = 9.78) compared to the 1st assessment (*M* = 80.41, *SEM* = 12.72). Overall, ratings were higher under placebo (*M* = 67.21, *SEM* = 16.57) than control (*M* = 42.14, *SEM* = 13.06). Together these results suggest absence of placebo effects in ratings of sinusoidal stimulation and/or masking by time effects.

In contrast, pain ratings during half-sinusoidal stimulation were overall lower under placebo (*M* = 45.78, *SEM* = 18.20) than control (*M* = 58.87, *SEM* = 21.24), consistent with placebo hypoalgesia. These ratings showed only a small decrease from the 1st (*M* = 52.45, *SEM* = 21.14) to the 2nd assessment (*M* = 49.70, *SEM* = 16.03). The absence of placebo effects in the ratings during sinusoidal stimulation may reflect a potential contrast or mismatch effect, where large discrepancies between sensory input and analgesia expectations may attenuate or even reverse placebo responses [8,9]. In contrast, half-sinusoidal stimulation, typically perceived as less intense, may have provided a more compatible stimulus range, resulting in placebo-consistent effects.

The present findings provide novel proof-of-principle evidence that expectations can modulate peripheral nociceptor activity. Given the small sample size and variability across protocols, results have to be interpreted cautiously. However, variability in the present results mirrors the complexity of peripheral nociceptive processing. Methodological aspects that could be optimized in future studies could be shorter test protocols to reduce time effects and the use of paradigms that allow rapid switching between placebo and control conditions to facilitating reinstatement of expectations, for example, using sham transcutaneous electrical nerve stimulation (TENS) as placebo [10,11]. Larger samples and refined designs are required to determine under which conditions cognitive states can reliably influence which peripheral nociceptor functions.

## METHODS

### Participants

Eight healthy volunteers were recruited for the study. Participants reported no history of neurological, dermatological, or other long-term medical disorders and did not take any regular or acute medication 24 h prior to their participation. Participants were recruited at the Heinrich Heine University Düsseldorf and at the RWTH Aachen University through posted announcements and during academic lectures.

For two participants the experiment was discontinued and any data were excluded, because no stable recordings from single C-fibers could be obtained during microneurography. The final dataset consisted of six participants (*M* = 25.33, *SD* = 5.82), with four female participants. All participants provided written informed consent prior to participation. At the end of the study, participants were fully debriefed about the purpose of the study and informed that both gels applied during the experiment were identical and contained no active pharmacological ingredients. The study was approved by the local ethics committee of the Uniklinik Aachen (EK 143-21 amendment) and conducted in accordance with the Declaration of Helsinki.

### Experimental Design

The study employed a within-subject, randomized crossover design. Each participant underwent both experimental conditions, a placebo condition and a control condition, within a single experimental session with a total duration of approximately 4 h. The order of conditions was counterbalanced across participants, i.e. three participants received the placebo condition first, followed by the control condition, while the remaining three participants underwent the conditions in opposite order. As microneurographic recordings yield a variable number of fibers per participant that can be analyzed, it was not feasible to have a counterbalanced order of placebo and control condition at fiber level.

Microneurographic recordings were performed across both experimental conditions (placebo and control). The aim was to assess every identified C-fiber under both experimental conditions. However, due to the technical characteristics of microneurography, including occasional loss of single-unit isolation or changes in recording stability over time, not all fibers could be successfully tested in all protocols and under both conditions. Consequently, sample sizes vary across microneurographic measures and conditions.

Participants were blinded to the experimental manipulation. They were instructed that one gel had analgesic properties and that this gel was compared to a control gel without any active ingredient. The experimenter was not blinded to condition, because maintaining the verbal instructions and conditioning procedures required knowledge of the experimental condition.

### Placebo paradigm

A well-established placebo paradigm [12,13] was employed, in which an inert gel was applied under two experimental conditions:

1. a *placebo* condition, in which expectations of pain relief were induced through verbal suggestion and classical conditioning, and
2. a *control* condition, in which the identical gel was applied with neutral instructions without suggestion of any pain-modulatory effects and without conditioning.

#### Inert gel and cover story

The gel (approx. 2 ml per application), applied in both conditions, contained 1,2-propylene glycol (10%), carbopol 974P (1%), trometamol (1%), disodium EDTA (0.1%), and water (87.9%) and did not include any active pharmacological ingredients.

As part of the cover story, participants were informed in the placebo condition that the gel contained a novel analgesic compound designed to selectively block a specific sodium channel, thereby reducing pain perception mediated by C-fibers while preserving tactile sensitivity. Participants were further instructed that short-lasting analgesic effects had been demonstrated in previous studies, but potential effects on peripheral C-fiber activity had not yet been investigated. In the control condition, the identical gel was used and described as inert and without any analgesic properties to serve as a control condition for the expected pain inhibitory effects in the experimental condition.

Both gels were applied in an identical manner across conditions and reapplied approximately every 20 min throughout the experiment to maintain the plausibility of the treatment effects for the cover story. Separate, visibly distinct containers for the gel were used for the placebo and control conditions to support the credibility of the cover story.

#### Verbal suggestion

Verbal suggestions were provided in both conditions. In the placebo condition, participants were informed that the gel has analgesic effects. In the control condition, participants were informed that the applied gel had no active ingrediencies and thus no analgesic effects to serve as a neutral comparison. Verbal suggestions were repeated during the experiment to maintain the expectations over time.

#### Conditioning procedure

At the beginning of the experimental session, a classical conditioning procedure was conducted to associate expectations of pain relief with the placebo (gel) through learning. The purpose of the conditioning procedure was to induce and enhance expectations of reduced pain under placebo treatment. During this procedure, the placebo gel was repeatedly paired with a reduced intensity of the applied electrical test stimuli in the placebo condition. In the control trials, electrical stimuli were presented with unaltered (higher) intensity to serve as a non-conditioned reference condition. Due to the requirements of microneurographic recordings, the conditioning procedure was performed on the foot contralateral to the microneurographic recording site. Two test areas on this foot were selected. For the conditioning procedure, sinusoidal electrical stimulation (12 s duration) was applied using handheld transcutaneous electrodes in form of two parallel platinum threads with contact length of 3mm in each area as used before [14]. Stimulation intensities were individually calibrated for each area by gradually increasing the current in predefined steps (0.02–0.5 mA) aiming at a target rating of 170 on a visual analogue scale (VAS) from 0–200, with 100 indicating the pain threshold.

One test area on the food was assigned randomly to the placebo condition and the other to the control condition. Following application, the gel was left on the skin to absorb for 5 min. The conditioning procedure consisted of 12 stimulation trials (six per condition) in total, presented in fixed order (three control trials followed by three placebo trials, repeated once). During placebo trials, the stimulation intensity was reduced to evoke sensations close to the individual pain threshold (approx. 100 on the VAS). Participants were not informed about the conditioning procedure, rather they were instructed that stimulation intensity was identical across trials and that any perceived changes in intensity would reflect the effect of the applied gel. Participants were further informed that this procedure was to assess how strongly the analgesic gel reduced pain perception on the level of the individual experience.

#### Manipulation check of analgesic expectation

To assess whether the conditioning procedure successfully induced expectations of pain relief, participants rated perceived pain-inhibitory effects of the placebo gel immediately after finishing the conditioning procedure. Ratings were provided on a numeric scale ranging from 0 (“not at all”) to 10 (“extremely”).

#### Gel application during microneurography

During the microneurographic recordings, the gel was applied in both the placebo and the control condition to the receptive field of the recorded fibers around the electrodes for electrical stimulation during microneurography (see below). Gel application, combined with the verbal suggestions, were repeated approximately every 20 min to maintain the plausibility of sustained treatment effects throughout the session.

### Microneurography recordings

To record single action potentials from single nerve fibers in humans, the electrophysiological technique Microneurography was performed following well-established procedures [7,15,16]. Recordings were obtained from cutaneous C-fiber fascicles of the superficial peroneal nerve. Participants were positioned comfortably in a supine position on a padded examination bed, with both legs supported to minimize movement throughout the experiment.

Single-nerve fiber recordings were obtained using tungsten microelectrodes (Frederick-Haer, Bowdoinham, ME, USA) inserted percutaneously near the nerve trunk and then moved towards the nerve until the uninsulated needle tip was located within a fascicle containing several nerve fibers (axons). In this position, mechanically evoked nerve fiber discharges were recorded, amplified and rendered audible through a loudspeaker to identify the receptive field. By gently scratching the skin with the fingers and listening to the characteristic sound of the mechanically evoked nerve fiber discharges, the receptive field of the respective fascicle was determined. Once a stable recording position was achieved, innervation territories of individual C-fibers were localized by transcutaneous electrical stimulation of the skin using a pointed search electrode (0.5 mm diameter). If action potentials were evoked by this electrical stimulation, C-fiber action potentials were identified in the recording by their slow conduction velocity (< 2 m/s) and individually stable response latency to repetitive electrical pulses with low frequency (0.25 Hz).

For repetitive 0.25 Hz stimulation throughout the experiment, a pair of fine needle electrodes (Frederick-Haer) was inserted into the identified receptive field of one or several C-fibers on the dorsum of the foot. Electrical stimuli were delivered using a constant current stimulator (Digitimer DS7A, Digitimer Ltd, Welwyn Garden City, Hertfordshire, United Kingdom). For technical reasons, for sinusoidal and half-sinusoidal stimulation protocols electrical stimuli were delivered using a different constant current stimulator (Digitimer DS5, Digitimer Ltd., Welwyn Garden City, Hertdordshire).

#### Experimental procedure and electrical stimulation protocols

Following completion of the placebo conditioning procedure, microneurographic recordings and stimulation protocols were conducted in a fixed order to ensure comparability across participants and fibers. At the beginning of each condition (placebo, control), the respective gel was applied around the electrodes for electrical stimulation, and verbal suggestions consistent with the experimental condition were provided. We used different, common microneurography protocols to compare different indicators of peripheral fiber activation and functionality between placebo and control condition.

Within each condition, the following microneurographic protocols were applied in the following order:

#### (1) Electrical activation threshold

To first determine the electrical activation threshold of single C-fibers, intracutaneous electrical stimulation delivered via the needle electrodes inserted into the receptive field was applied. Thresholds were assessed by reducing the current intensity to a level that reliably evokes action potentials. The lowest current that reliably evoked action potentials was defined as the “threshold intensity”. Electrical stimulation was thereafter performed at double the threshold for the highest threshold nerve fiber. In case this would be too painful for participants, 1.5 threshold was used.

#### (2) Electrical identification/low-frequency stimulation

To assess activity-dependent conduction velocity slowing (ADS) during low-frequency electrical stimulation, a series of electrical pulses was delivered directly to the skin via the needle electrodes following a two-minute pause. Fibers were stimulated with 0.5 ms square-wave pulses at progressively increasing frequencies: 1/8 Hz (20 pulses), 1/4 Hz (20 pulses), and 1/2 Hz (30 pulses). Conduction latency was then allowed to recover at 1/4 Hz for an additional 10 pulses. This stimulation protocol was used to quantify ADS as a biophysical measure of fiber excitability and to distinguish between different fiber types, including mechano-sensitive (CM) and mechano-insensitive (CMi) C-fibers. ([17]; see below “C-fiber classification”).

#### (3) Mechanical thresholds

To assess the mechanical sensitivity of the recorded C-fibers, repetitive mechanical stimuli were applied to the receptive fields on the dorsum of the foot using a stiff von Frey filament (Stoelting, Chicago, IL, USA). Stimulation was performed within an area of approximately 3 cm surrounding the needle electrodes. Activation was defined as a sudden increase in conduction latency, followed by a gradual, stepwise normalization of this latency. This protocol was used for fiber classification (see below “C-fiber classification”).

#### (4) Velocity recovery cycles (“old double pulse”)

Alongside continuous 0.25 Hz stimulation, an additional rectangular electrical pulse is administered approx. every tenth pulses at various intervals preceding the regular pulse [18]. The interval between the continuous stimulation and the regular pulse ranged from 2000ms to 20ms. Changes in conduction latency of the second action potential were analyzed as a function of interstimulus intervals to assess early post-spike excitability changes, including phases of reduced or increased excitability. In the phases of early (20-50ms interpulse intervals) and late (>250ms) subnormality, the second action potential is conducted slower. With a certain amount of preceding activity, a so called “supernormal phase” occurs. During this period around 30-250 ms after the action potential, the second action potential is conducted faster than the first one. Only in CMi the preceding activity caused by 0.25 Hz electrical stimulation is sufficient to cause such a supernormal period. In other fibers, higher frequencies would be necessary. This faster conduction leads to a clustering of action potentials and dramatically increased frequencies arriving at the spinal cord. A reduction of the supernormal phase would lead to a dispersion of action potentials and lower frequencies arriving at the spinal cord [19]. Taken together the recovery cycle can be used as indirect indicator of axonal excitability and axonal signal processing leading to modulated discharge patterns arriving at the spinal cord.

#### (5) Sinusoidal Stimulation

Sinusoidal electrical stimulation is used because it effectively activates CMi-fibers, but not A-fibers including nociceptive A-delta fibers [14]. By contrast, rectangular stimuli activate CMi-fibers only with high intensities, which due to spatial summation, are often too painful for the volunteers. Sinusoidal stimuli (4Hz) were applied for a duration of 12 s. Stimulation intensity was increased stepwise across trials, ranging from 0.025 to 0.4 mA. This protocol was used to assess fiber responses to sustained rhythmic stimulation and to determine stimulation threshold and latency changes under dynamic stimulation conditions. The stimuli used in this protocol elicited a distinct perception that allowed ratings of perceived intensity. Following each stimulation trial, participants therefore rated their perceived pain intensity using the same VAS as before (from 0 to 200, with 100 indicating the pain threshold).

#### (6) Half-Sinusoidal Stimulation

Half-sinusoidal electrical pulses with a duration of 0.5 s were applied to assess responses to brief, discrete stimuli [20,21]. Half sine shaped electrical stimuli evoke a short burst in the activated fibers. Thus, this stimulus tests for the ability of a nerve fiber to discharge a high frequent burst. Four predefined stimulation intensities were used (0.1, 0.2, 0.6, and 1.0 mA). Each intensity was presented once per condition, depending on recording stability. This protocol also used stimuli that elicited a distinct perception, allowing ratings of perceived intensity. After each stimulation, participants rated their perceived pain intensity on the same VAS as before.

#### C-fiber Classification

According to their mechanical response (see “(3) Mechanical thresholds”) and electrophysiological properties (see “(2) Electrical identification”), we classified C-fibers into mechano-sensitive fibers (CM) and mechano-insensitive fibers (CMi) as performed before [15]. C-fibers with an activity-dependent conduction velocity slowing (ADS) of less than 5% of their initial latency to an electrical stimulation protocol with rising frequencies, a normalization of latency thereafter of more than 44% within 40 seconds and a response to 22 g von Frey stimulation were classified as mechano-sensitive fibers (CM). C-fibers with more than 5% ADS and less than 44% recovery and no response to mechanical stimuli were classified as mechano-insensitive nociceptors (CMi).

### Data Processing

Microneurography data were amplified, band-pass filtered, and digitized for online monitoring using DAPSYS 8 software (Brian Turnquist, Bethel University, St. Paul, Minnesota). Offline analyses were performed using DAPSYS and Microsoft Excel. Ratings of perceived stimulus intensities were given verbally and recorded manually by the experimenter during the experiment.

### Data Analysis

#### Microneurography

Conduction latency of action potentials was determined as the time between time of the stimulus and the occurrence of the action potential. To account for baseline differences between fibers, latency measures were normalized to each fiber’s unconditioned baseline conduction latency after a 2-minute break without any stimulation within the respective protocol. For recovery cycle analyses, changes in conduction latency of the second action potential of the double pulses were assessed. Recovery cycle parameters were analyzed descriptively by comparing latency shifts across interpulse intervals between placebo and control conditions.

Electrical and mechanical thresholds as well as latency measures obtained during the different stimulation protocols were compared descriptively between experimental conditions (placebo vs. control) and between first and second assessments to examine potential condition-related and time-dependent effects.

### Behavioral data

#### Sinusoidal stimulation

For sinusoidal stimulation, stimulus intensity ratings were averaged for each participant for the placebo and control conditions separately across all stimulation intensities (see “(5) Sinusoidal stimulation”). These mean ratings were then averaged across participants to obtain group means for the placebo and control condition, reported with the standard error of the mean (SEM). To examine time effects, ratings were averaged for each participant for the first and second assessments separately across all stimulation intensities that were applied in both assessments, independent of condition. These means were then averaged across participants and the group means for the first and second assessments are reported with the SEM.

#### Half-sinusoidal stimulation

For half-sinusoidal stimulation, stimulus intensity ratings were averaged within each participant across the four stimulation intensities for the placebo and control condition separately (see “(6) Half sinusoidal stimulation”). These mean ratings were then averaged across participants to obtain group means for each condition, which are reported with SEM.

To assess time effects, ratings were averaged within each participant across the four stimulation intensities for the first and second assessments separately. These means were then averaged across participants and the group means for the first and second assessment are reported with the SEM.

Given the exploratory and proof-of-principle nature of the study and the limited number of participants and recorded fibers all analyses were descriptive. No inferential statistical testing was performed. Results are reported as means with standard error of the mean (SEM) where applicable.

## ACKNOWLEDGEMENTS

The authors thank Danxia Bao for her technical support for microneurography and Lara Marcia Dittmann for her support with illustrations for the graphical abstract. B.N. is supported by DFG NA970/6-2, DFG NA970/7-1, and DFG NA970/9-2.

## GRAPHICAL ABSTRACT

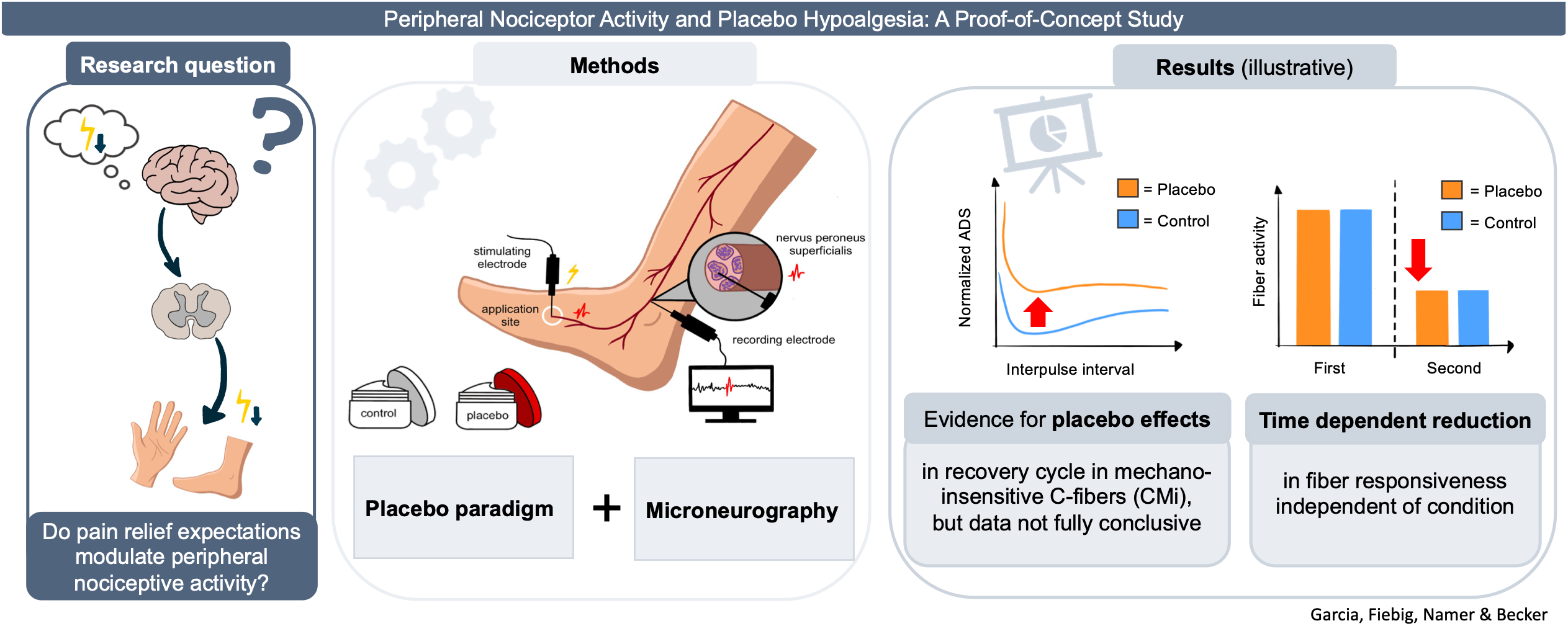

